# Aliphatic residues contribute significantly to the phase separation of TDP-43 C-terminal domain

**DOI:** 10.1101/2022.11.10.516004

**Authors:** Priyesh Mohanty, Jayakrishna Shenoy, Azamat Rizuan, José F Mercado Ortiz, Nicolas L. Fawzi, Jeetain Mittal

## Abstract

TAR DNA binding protein 43 (TDP-43) is involved in key processes in RNA metabolism such as splicing, stability and transcription. TDP-43 dysfunction is frequently implicated in many neurodegenerative diseases, including amyotrophic lateral sclerosis (ALS) and fronto-temporal dementia (FTD). The prion-like, disordered C-terminal domain (CTD) of TDP-43 is aggregation-prone and harbors the majority (~90%) of all ALS-related mutations. Recent studies have established that TDP-43 CTD can undergo liquid-liquid phase separation (LLPS) in isolation and is important for phase separation (PS) of the full-length protein under physiological conditions. While a short conserved helical region (CR, spanning residues 319-341) promotes oligomerization and is essential for LLPS, aromatic residues in the flanking disordered regions (IDR1/2) have also been found to play a critical role in PS and aggregation. However, TDP-43 CTD has a distinct sequence composition compared with other phase separating proteins, including many aliphatic residues. These residues have been suggested to modulate the apparent viscosity of the resulting phases, but their direct contribution to phase separation has been relatively ignored. Here, we utilized a multiscale simulation and experimental approach to assess the residue-level determinants of TDP-43 CTD phase separation. Single chain and condensed phase simulations performed at the atomistic and coarse-grained level respectively, identified the importance of aromatic residues (previously established) while also suggesting an essential role for aliphatic methionine residues in LLPS. *In vitro* experiments confirmed the role of phenylalanine, methionine, and leucine (but not alanine) residues in driving the phase separation of CTD, which have not been previously considered essential for describing the molecular grammar of PS. Finally, NMR experiments also showed that phenylalanine residues in the disordered flanking regions and methionine residues both within and outside the CR contribute important contacts to CTD interactions. Broadly, our work highlights the importance of non-alanine aliphatic residues such as methionine and leucine, and potentially valine and isoleucine, in determining the LLPS propensity, expanding the molecular grammar of protein phase separation to include critical contributions from aliphatic residues.

## Introduction

Membraneless organelles, also referred to as biomolecular condensates, organize cellular biochemistry by sequestering functionally-related macromolecular components from the bulk cellular compartment environment (Banani et al., 2017; Sabari et al., 2020; Schuster et al., 2021). Experimental studies indicate that many of these organelles form via phase separation to create distinct subcompartments within either the nucleus or cytoplasm (Hyman et al., 2014). These organelles include nucleoli, Cajal bodies, stress granules (SGs), and other ribonucleoprotein (RNP) granules (Rhine et al., 2020). SGs, which primarily serve to sequester and halt the translation of mRNAs (Protter and Parker, 2016), recruit many distinct RNA-binding proteins (RBPs) and can form dynamically in response to physiological stress. In recent years, RBPs containing disordered, prion-like domains (eg. FUS, hnRNPA2, hnRNPA1, TDP-43) have received considerable attention due to their involvement in stress granule assembly and maintenance (Harrison and Shorter, 2017). The prion-like, disordered regions (PrLDs) are typically enriched in polar residues (Ser, Thr, Asn and Gln) and can undergo liquid-liquid phase separation (LLPS) in cells and as purified proteins *in vitro* via weak, multivalent interactions (Mittag and Parker, 2018). Phase-separated droplets of prion-like domains can also mature to form fibril-like structures (Babinchak and Surewicz, 2020) which may exert a toxic cellular effect.

One constituent of stress granules and other RNP granules that has received significant attention is TAR DNA binding protein 43 (TDP-43), a primarily nuclear RBP that plays an essential role in the regulation of mRNA splicing (Cohen et al., 2011). Mislocalization of TDP-43 to the cytoplasm due to chronic stress and/or in combination with mutations that disrupt its structure and interactions gives rise to the formation of ubiquitinated neuronal inclusions (Neumann et al., 2006), a hallmark of amyotrophic lateral sclerosis (ALS). TDP-43 has a multi-domain architecture consisting of an N-terminal folded domain that can form linear globular chains (Afroz et al., 2017; Wang et al., 2018a), two RNA recognition motifs (RRMs) that together bind RNA (Lukavsky et al., 2013), and a prion-like C-terminal domain (CTD) which is predominantly disordered (Conicella et al., 2016; Lim et al., 2016). As a purified protein *in vitro* conditions, the C-terminal domain readily phase separates in the presence of physiological concentrations of salt or RNA (Conicella et al., 2016) which is enhanced at low temperatures (Li et al., 2018a). Within the CTD, there is an evolutionarily-conserved, hydrophobic sequence that takes on α-helical structure (Conicella et al., 2016; Lim et al., 2016) and is flanked by disordered regions - IDR1, Q/N, and IDR2 (Fig. S1). Homo-oligomerization through this conserved region (CR) makes a large contribution to phase separation of the CTD; changes in helical stability through either ALS-related or designed mutations (Conicella et al., 2020; Conicella et al., 2016; Li et al., 2018a; Schmidt and Rohatgi, 2016) were shown to tune phase separation. In addition to CR, aromatic residues in the flanking regions (IDR1/2) were also shown to be critical for LLPS (Li et al., 2018b; Schmidt et al., 2019) and fibrillation of CTD (Pantoja-Uceda et al., 2021).

A detailed bioinformatic analysis revealed the sequence conservation in IDR1/2 and the high conservation in the spacing of aromatic and hydrophobic residues (Schmidt et al., 2019). As shown in Figure 1, TDP-43 CTD contains 8 phenylalanine (6 of them regularly spaced within the disordered flanks), 3 tryptophan and 1 tyrosine. Both *in vitro* and *in vivo* experiments (Li et al., 2018a; Li et al., 2018b; Schmidt et al., 2019) point toward the essential nature of aromatic residues (F, W, Y) in driving the phase separation of TDP-43. While arginine-mediated interactions were found to be important for the condensed dynamics, neither electrostatics nor arginine-mediated interactions within the CTD were able to inhibit the LLPS of the TDP-43 reporter construct (Schmidt et al., 2019). While the importance of aromatic and charged residues in phase separation of TDP-43 CTD and other prion-like domains is well-established (Bremer et al., 2022; Martin et al., 2020; Schuster et al., 2020; Wang et al., 2018b), the relative contributions of other residues (e.g., aliphatic residues) to LLPS is less understood – previously they have been viewed as playing a role in tuning the dynamics and solidification or aggregation, but not playing a major role in driving phase separation (Schmidt et al., 2019). It is worth considering the abundance of aliphatic hydrophobic residues within and outside of the CR - 7 alanine, 5 methionine, and 2 leucine in TDP-43 CTD each in both the CR and the flanking IDRs. In this study, we address whether these aliphatic residues are important for determining the PS propensity of TDP-43 CTD. Furthermore, how residue types outside the CR affect the TDP-43 CTD self-assembly and how their contribution differs from the critical residue types within the CR is interrogated.

**Figure 1.**
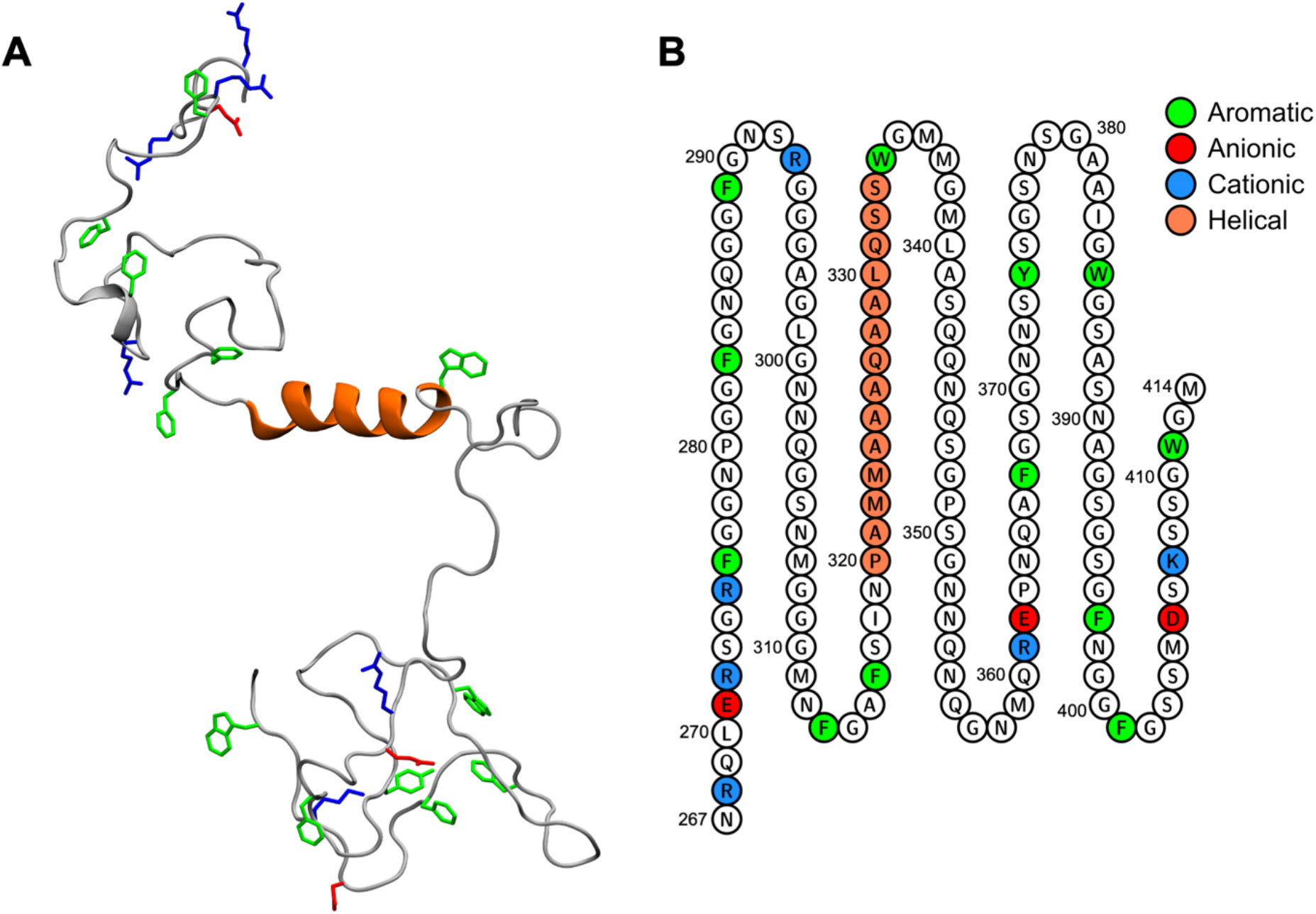
Sequence composition of the TDP-43 C-terminal domain (CTD) **A**. Cartoon representation of the C-terminal domain of TDP-43 that contains aromatic and charged residues outside the conserved region (CR, spanning residue 319-341) enriched in aliphatic residues **B**. TDP-43 CTD (aa: 267-414) sequence. Letters in green, red, and blue circles are aromatic, anionic, and cationic residues, respectively. The residues within CR of TDP-43 CTD are shown as orange circles.

Significant advancements in the accuracy of enhanced sampling algorithms (Barducci et al., 2015) and atomistic force fields for simulating intrinsically disordered proteins (Robustelli et al., 2018; Shea et al., 2021) allow for an in-depth characterization of the conformational landscape of intrinsically disordered proteins (Bernetti et al., 2017; Lindorff-Larsen et al., 2012; Shrestha et al., 2019; Zerze et al., 2015), which complements insights obtained from biophysical experiments such as NMR, SAXS, and FRET (LeBlanc et al., 2018; Sibille and Bernado, 2012). Further, coarse-grained simulations parameterized based on amino acid hydrophobicity scales (Dignon et al., 2018b; Regy et al., 2021) have provided valuable insights into the relationship between sequence and phase behavior of intrinsically disordered proteins. Here, we employed a multiscale simulation strategy based on atomistic and coarse-grained simulations coupled with *in vitro* experiments and high-resolution Nuclear Magnetic Resonance (NMR) spectroscopic data to probe in detail the role of residue types on TDP-43 phase separation. Looking ahead, our results expand the atomically detailed picture of TDP-43 CTD phase separation that, in turn, helps to understand its function and aggregate pathology in neurodegenerative disease. Our results also highlight the unexpected and underappreciated role of residues such as methionine and leucine in protein PS.

## Results

### Single chain simulation accurately captures the conformational ensembles of TDP-43 C-terminal domain

The degree of conformational collapse of disordered proteins at dilute concentrations positively correlates with their ability to undergo LLPS at higher concentrations (Dignon et al., 2018a; Lin and Chan, 2017). Previously, we performed extensive characterization of small disordered fragments (< 50 amino acids) belonging to RBPs such as LAF-1 (Schuster et al., 2020), FUS (Murthy et al., 2019), and hnRNPA2 (Ryan et al., 2018) at the single chain level. Because these phase-separating disordered domains contain repetitive sequence regions with low complexity, the analysis of intramolecular contacts observed in single-chain simulations mimicking the dispersed phase provided insights into intermolecular interactions implicated in phase separation. Recently, single chain simulations of the longer Tau - K18 repeat (130 amino acids) were used to understand the origin of its non-linear, temperature-dependent phase behavior (Dong et al., 2021). Based on the above findings, we performed single chain simulations of CTD (aa: 267-414) to characterize its intramolecular interactions which maybe relevant for phase separation.

Since TDP-43 contains a conserved helical region (aa: 319-341), it is important to ensure that the structural propensity of this region is accurately captured in simulation. In our earlier studies (Conicella et al., 2020; Conicella et al., 2016), single chain simulations of the TDP-43 CTD fragment (aa: 310-350) performed using parallel tempering simulations in the well-tempered ensemble (PT-WTE) (Deighan et al., 2012) faithfully captured the helical propensity of CR observed by NMR. Here, we performed PT-WTE simulation of the entire CTD (aa: 267-414) and first characterized its conformational properties. After establishing sufficient convergence (see Methods, Fig S2), the CTD ensemble at 300 K was compared to experimentally observed NMR chemical shifts and secondary structure propensities (Conicella et al., 2016). For the simulated ensemble, backbone ^13^C_α_ and ^13^C_β_ chemical shifts, which provide residue-specific information on the secondary structure at each position, were predicted from the simulation ensemble using the SPARTA+ algorithm (Shen and Bax, 2010). RMSDs of chemical shifts with respect to experiment were less than 0.5 ppm which is within the prediction error and indicates good agreement (Fig. S3A) similar to what we showed before for a subsegment of TDP-43 C-terminal domain (Conicella et al., 2016). Per-residue secondary structure populations computed from simulation using DSSP (Kabsch and Sander, 1983) indicate helical propensity for residues in CR (Fig. S4). The helical populations in the ensemble appear to be in reasonable agreement with those computed based on experiment chemical shifts using *δ*2D (Camilloni et al., 2012). *δ*2D predictions depend on the direct mapping of chemical shifts to secondary structure populations and utilize a training data set which contains both globular and disordered proteins. The accuracy of such predictors is, however, limited by the quality of the training data set. A direct assessment of helical populations can be achieved by comparison of per-residue secondary chemical shift differences between simulation and experiment (Fig. 2A, S3B). In agreement with experiment, positive values of (>0.6 ppm) of the ^13^C shifts compared to those expected for a random coil reference (ΔδCα and ΔδCα - ΔδCβ shifts) were observed for CR residues (aa: 320-330 and 335-342) which indicates the presence of α-helical structures (Conicella et al., 2016; Lim et al., 2016). In conclusion, PT-WTE simulations generated an accurate ensemble of CTD, which is suitable for the analysis of intramolecular interactions relevant to its phase separation.

**Figure 2.**
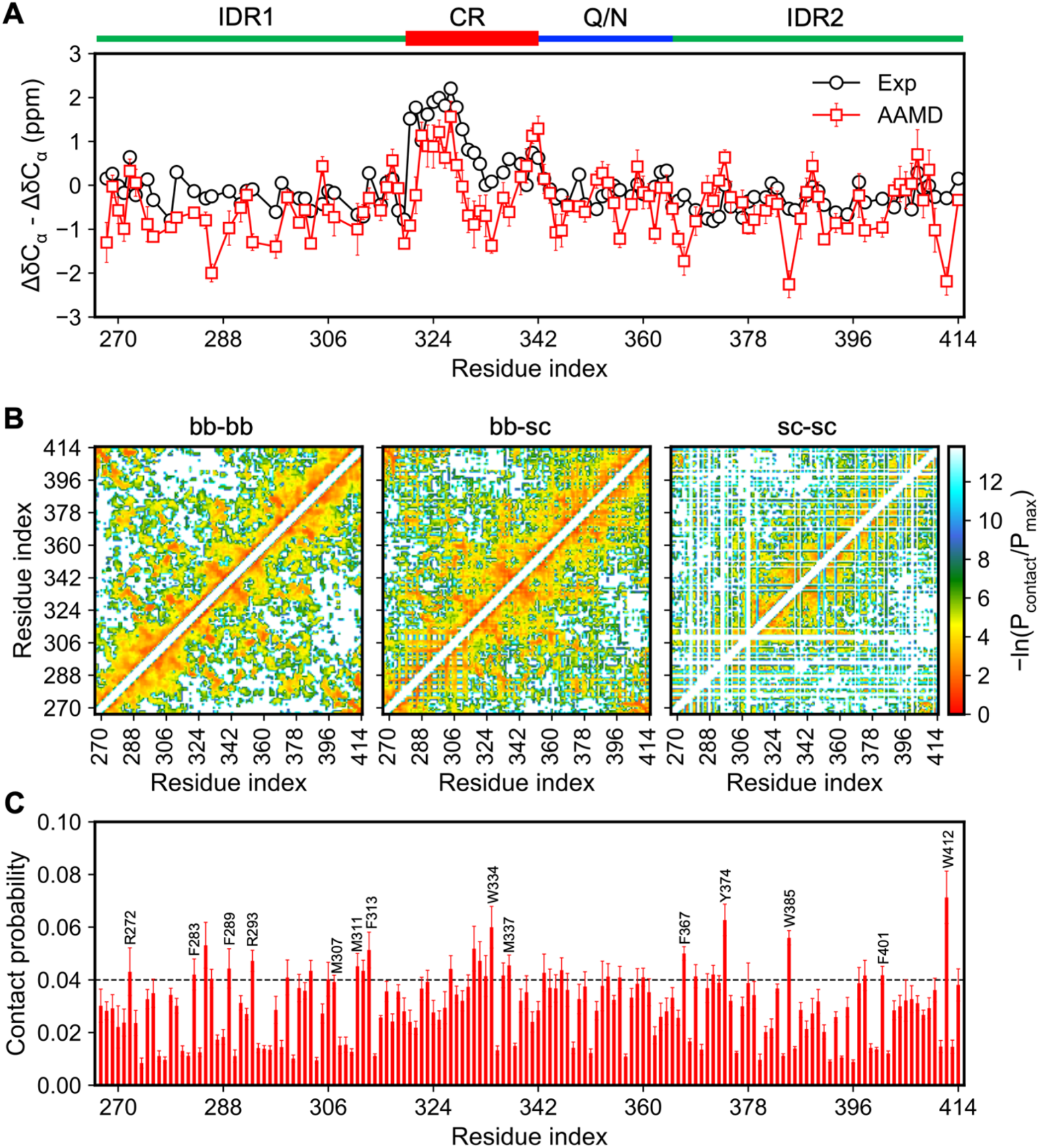
Single chain simulation of TDP-43 CTD at atomistic resolution captures the structural conformations and identifies important intramolecular interactions. **A**. Validation of CTD ensemble at 300 K based on NMR observables. Per residue, secondary chemical shifts from 300 K replicas show a good agreement with the experiment. **B**. 2D contact maps show van der Waals contacts as a function of residue index, classified based on backbone-backbone, backbone-sidechain and sidechain-sidechain interaction modes. **C**. Per-residue contact probabilities derived through summation of pairwise contact probabilities (bb-sc + sc-sc) at each residue position based on 2D contact maps shown in B. Block averaging was performed over 100 ns intervals to obtain the mean and standard errors associated with each residue. The dashed line corresponds to the mean+σ probability cutoff used to identify individual residues which exhibit high contact probabilities. Analysis of intramolecular contacts suggests that multiple residues within and outside the CR may contribute toward CTD phase separation.

### Single chain simulation identifies important contacts in TDP-43 C-terminal domain

To probe the contacts that mediate TDP-43 CTD interactions, we performed an in-depth analysis of pairwise residue contacts in the CTD ensemble generated by PT-WTE simulations. Two-dimensional contact maps of pairwise interactions based on residue index are shown in Figure 2B. The interactions are classified into backbone-backbone (bb-bb), backbone-sidechain (bb-sc), and sidechain-sidechain (sc-sc) interaction modes. We find that residues within CTD contribute to the pairwise contacts via all three interaction modes, although sc-sc interactions are sparser owing to the abundance of glycine with only hydrogen atom in the side chain. Next, position-specific interactions in the CTD sequence were also analyzed to identify residue positions that show high contact probabilities. To do so, we first derived per-residue contact probabilities for each interaction mode through summation of pairwise contact probabilities at each position based on 2D contact maps (Fig. 2C). To assess the significance of each interaction mode, we also computed per-residue interaction energies (Zeron et al., 2019) and analyzed their correlation with the corresponding contact probabilities. A strong positive correlation was observed for bb-sc and sc-sc interactions, while bb-bb interactions showed a weak correlation (Fig. S5). Hence bb-sc and sc-sc interaction modes dictate the overall energetics of residue-level interactions. The per-residue contact probabilities summed over bb-sc and sc-sc interaction modes are shown in Figure 2C.

Based on per-residue contact probabilities, aromatic residues located outside the CR (F283/289/313/367, W385/412 and Y361) and R272/293 within IDR1 showed high contact probabilities compared to one standard deviation above the mean (mean+σ) which is consistent with the established role of aromatic residues in driving phase separation of CTD (Li et al., 2018b) and currently understood molecular grammar of the disordered prion-like domain phase separation (Bremer et al., 2022; Wang et al., 2018b). Intriguingly, methionine residues (M307/311/337) within and outside of the CR also showed high contact probabilities.

To examine the combined effects by a particular residue type, we next calculated the total contacts formed by specific residue pairs by bb-sc and sc-sc atoms (Fig. S6). As expected, the enrichment of glycine (G) and polar residues (S, N, and Q) in CTD results in an abundance of contact pairs that involve these residue types. As previously suggested, those residues may involve in variety of interaction modes such as backbone sp^2^-π (Vernon et al., 2018) and hydrogen bonding (Murthy et al., 2019). However, other studies have suggested that these residue types may only play a minimal role in phase separation, only influencing the material properties of disordered prion-like domain condensates (Wang et al., 2018b). As discussed above, we observed that aromatic (W and F) residues engage in a substantial number of interactions by its backbone and sidechain atoms. Consistent with observations from per-residue contact probabilities (Fig. 2C), methionines are among residues with higher contact propensity. Collectively, our contact analysis suggests that along with aromatic and polar residues, the presence of methionine residues may further increase the multivalency of CTD and contribute favorably towards its phase separation.

### Coarse-grained simulations predict the importance of non-alanine aliphatic residues in TDP-43 CTD phase separation

The observed intramolecular contacts in the single-chain ensemble from atomistic simulations may also contribute to the inter-chain interactions that stabilize the protein condensed phase (Dignon et al., 2018a). Therefore, the roles of key residues in CTD phase separation identified from the atomistic simulations can be further probed by performing phase coexistence simulations. Since AA simulations are computationally inefficient in studying the thermodynamic phase behavior, coarse-grained (CG) models that replace the atomistic details with a single bead per amino acid can be used to reach the time and length scales to study the phase separation of IDRs (Dignon et al., 2018b). For that reason, we conducted phase coexistence simulations of TDP-43 CTD using a recently developed CG model (Regy et al., 2021) based on the Urry hydrophobicity scale (Urry et al., 1992) (see Materials and Methods). Because this CG model does not explicitly represent secondary structure propensity based on sequence, in these simulations residues 320-341 were kept rigid in α-helical conformation using rigid body constraint (Nguyen et al., 2011) to partially mimic the structure identified by experiments and atomistic simulations. The qualitative behavior of TDP-43 CTD single-chain properties from the CG simulations is consistent with the CTD ensemble at 300K (Fig. S7). Although single-chain CG simulations sample more extended CTD conformations, residue-type contact probabilities sampled from the CG simulations are in good agreement with the results of atomistic simulations described above (Fig. S7). The mean value of R_g_ (3.05 ± 0.01 nm) from the single-chain CG simulations is also comparable to the approximate R_g_ of monomeric TDP-43 CTD (2.8 nm) based on its diffusion coefficient (Conicella et al., 2020). These observations give us further confidence to use the CG model to faithfully study the CTD condensed phase. The slab geometry is utilized for CG simulations of phase coexistence that allowed efficient equilibration between the dilute and condensed phases (Fig. 3A). The intermolecular van der Waals (vdW) contact map of wild-type (WT) CTD (Fig. 3B) shows each subdomain contributes to overall interactions to differing degrees, while self-interaction within the CR is slightly favored. Both per-residue and residue-type intermolecular contacts in the condensed phase were found to be in excellent agreement with intramolecular contacts from single-chain atomistic simulations (Fig. 3C, 3B, S8). These observations established a correspondence between single chain and condensed phase interactions and allowed us to test the ability of mutant CTD variants (Fig. 3A, 3D) to form a stable condensed phase.

**Figure 3.**
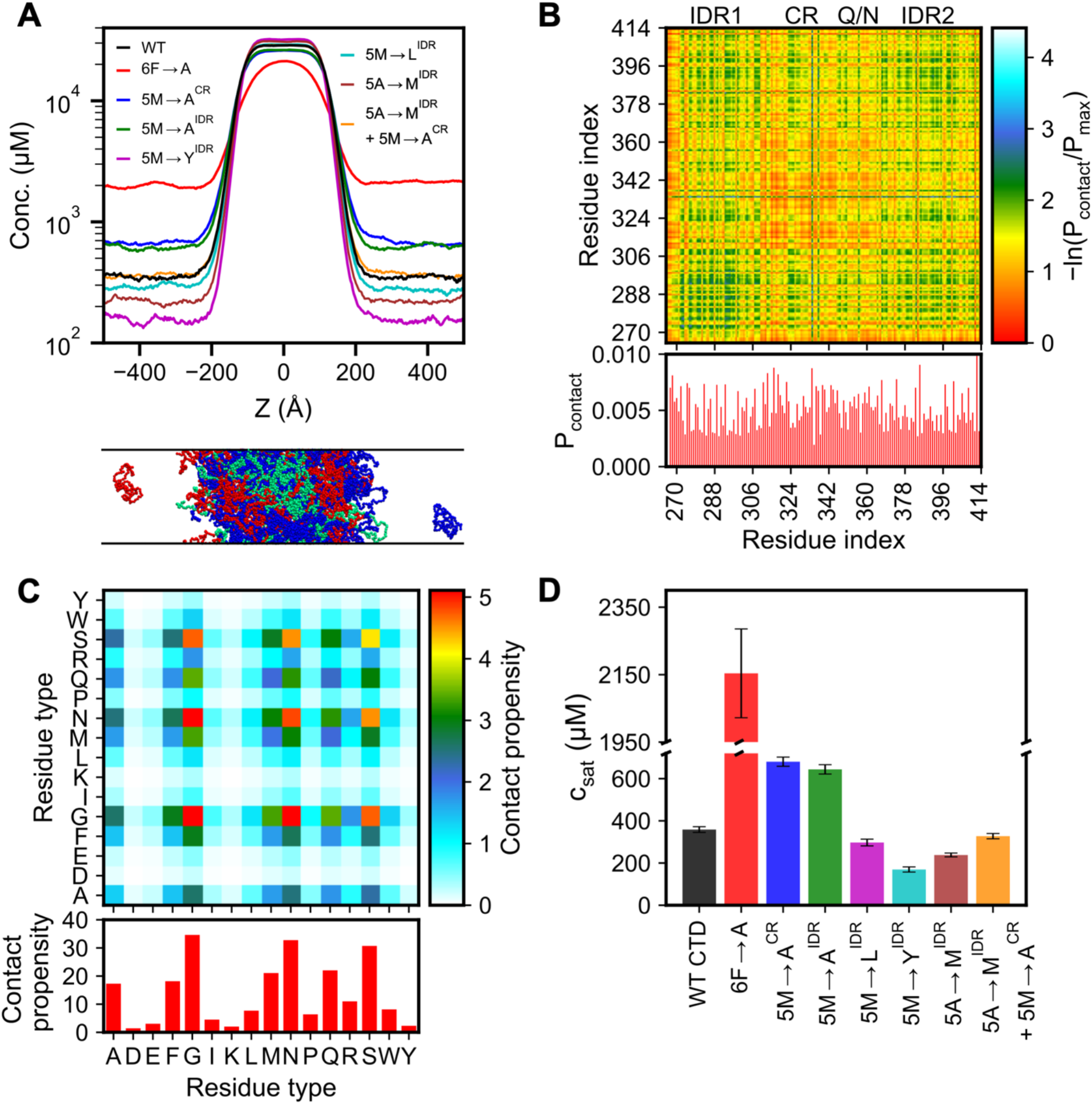
CG phase coexistence simulations indicate the importance of aromatic and bulky aliphatic residues in phase behavior of TDP-43 CTD. **A**. Snapshot of TDP-43 CTD WT slab configuration (100 chains) from CG simulation at 300K, where two phases coexist (bottom). The concentration profiles of TDP-43 CTD WT (black) and its variants along the z-dimension of the slab. (top) **B**. Pairwise intermolecular interactions (top) and contact probabilities per residue (bottom) in the condensed phase show the key residues involving phase separation. **C**. Pairwise interactions within the condensed phase as a function of residue type. **D**. Saturation concentration (c_sat_) for TDP-43 CTD WT and its variants computed based on concentration profiles in A.

Given the highest abundance aromatic and aliphatic residues are phenylalanine (8 in total) and methionine (10 in total), respectively, we first tested the impact of substitutions of these residue types on the stability of the condensed phase based on changes in the saturation concentration (c_sat_) relative to WT CTD (Fig. 3A). To do so, we designed mutant variants mutating these residues within and outside the CR independently (Table S1). It is important to mention that we carefully engineered those mutant variants to avoid the disruption of helicity present in the CR, which are critical for CTD self-assembly. The aromatic mutant – 6F→A (all phenylalanines to alanine, except F313, and F316, which are close to the CR) led to a >6-fold increase in c_sat,_ indicating significant destabilization of the condensate. Next, the effects of methionine residues in the CR (5M→A^CR^) and disordered flanking regions (5M→A^IDR^) were tested by substituting them with alanine, keeping in mind that alanine has higher experimental helical propensity than methionine (Moreau et al., 2009). Interestingly, mutational change in hydrophobicity by replacing methionine residues with alanine, both outside the CR (5M→ A^IDR^) and inside the CR (5M→ A^CR^), resulted in a lower phase separation propensity (~2-fold increase in c_sat_) which suggests the contribution for methionine residues for LLPS. Conversely, the substitution of methionines outside the CR with another aliphatic residue, leucine (5M→L^IDR^), shows a similar c_sat_ as WT, hinting that the contribution to phase separation may not be specific to methionine but that other large aliphatic (non-alanine) residues could substitute. Meanwhile, mutating all five methionine residues outside the CR region to tyrosine (5M→Y^IDR^) displayed substantial enhancement of phase separation (>3-fold increase in c_sat_). Thus, even if methionine (and leucine) are important contributors to phase separation, their relative contribution is lower than that of tyrosine, an aromatic residue, tyrosine, which is found to be crucial for LLPS of many IDPs, including the low complexity domains of FUS and hnRNPA1 (Bremer et al., 2022; Wang et al., 2018b). Importantly, these variants shed light on the residue-level determinants of CTD phase separation, which may be universal to other phase separating prion-like domains (PrLDs). Overall, the variation in c_sat_ observed for CTD mutants suggests that phenylalanines in the flanking disordered regions and methionine residues present within and outside the CR region significantly contribute to the phase separation of CTD.

Additional analysis of methionine residues via mutations using CG phase coexistence simulations suggests that methionines in both IDR1 (M307A, M311A) and IDR2 (M359A, M405A, M414A) affect the phase behavior of CTD while substituting all methionine to alanine (10M→A) predicted to drastically weaken the phase separation compared to methionine variants examined above (Fig. S9A). Positional mutational studies demonstrate that all individual methionine residues have contributions toward LLPS and collectively stabilize the condensed phase of TDP-43 CTD (Fig. S9B). These computational results further support the importance of methionine residues, which have not been previously recognized as essential for LLPS.

### Phenylalanine and methionine residues are crucial in TDP-43 CTD phase separation *in vitro*

To validate the observed trend in c_sat_ for WT CTD and its mutants observed in CG phase coexistence simulations, we tested the effect of these mutations on TDP-43 CTD phase separation *in vitro* by microscopy and droplet sedimentation assays (Fig. 4, Materials and Methods). Droplet sedimentation assays provide a quantitative assessment of phase separation, wherein the concentration remaining in the supernatant after centrifugation of the suspended droplets provides a measure of the saturation concentration (c_sat_) (Mackenzie et al., 2017) for the WT and variants that do not undergo rapid conversion into static aggregates. Under the given buffer conditions, phase separation occurs if the protein concentration is equal to or above its c_sat_.

**Figure 4:**
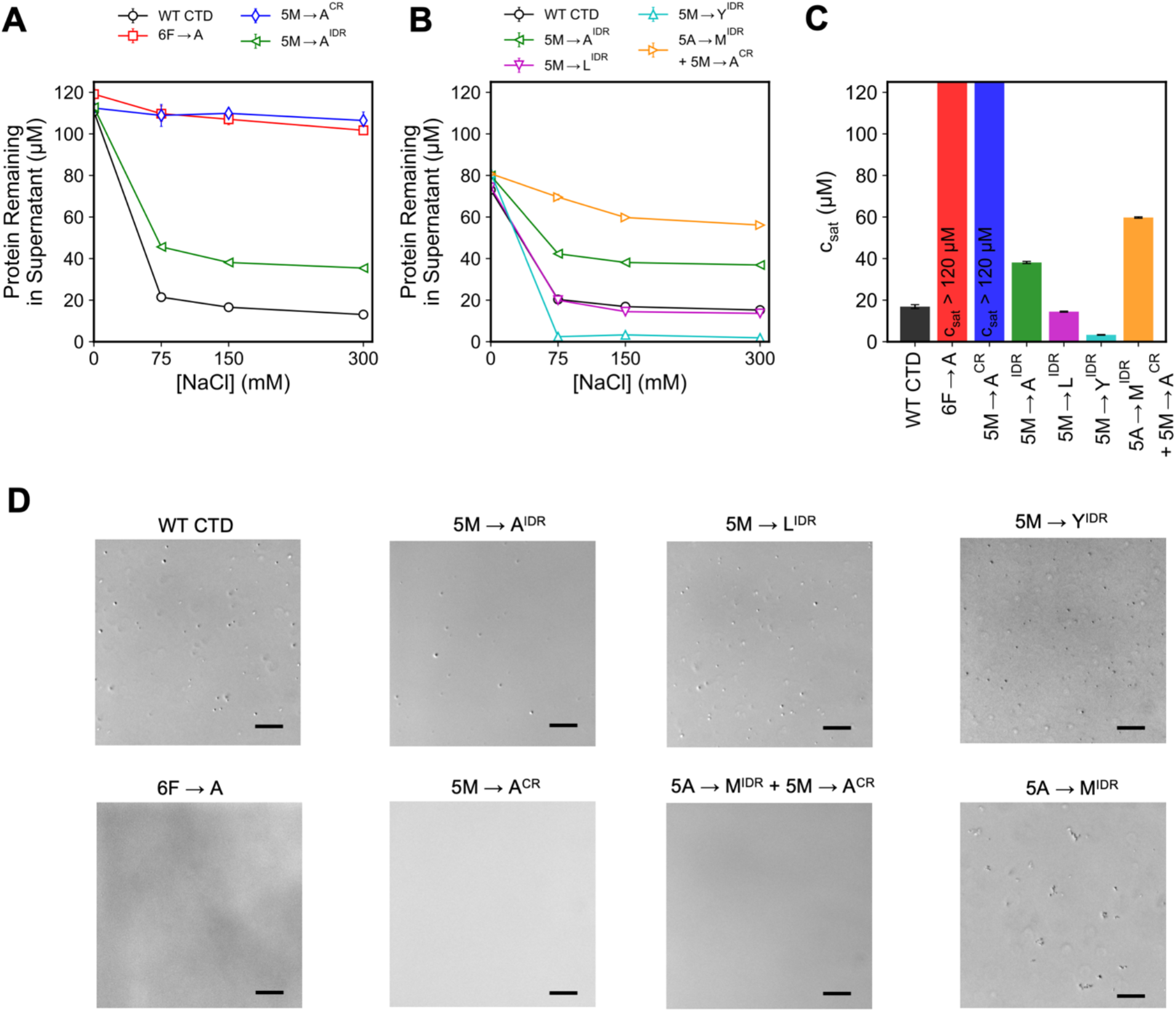
Phenylalanine and methionine residues are crucial in TDP-43 CTD phase separation *in vitro*. **A-B**. Quantification of phase separation assay showing the protein remaining in supernatant after phase separation for WT and its variants measured from 0 to 300 mM NaCl. Standard deviations of 3 replicates are represented as error bars. **C**. Saturation concentration (csat) for TDP-43 CTD WT and its variants are determined by measuring supernatant concentration at 300 mM NaCl **D**. DIC micrographs for WT TDP-43 CTD (20 μM) and designed variants at 300 mM salt. Scale bar, 20 μm.

WT CTD did not undergo phase separation in the absence of salt and showed an enhanced propensity for droplet formation upon increasing salt concentration (Fig. 4A), as shown previously (Conicella et al., 2020; Conicella et al., 2016). In contrast, 6F→A mutation impaired droplet formation up to 120 μM protein concentration across all salt concentrations (Fig. 4A). Similarly, 5M→A^CR^ variant inhibits phase separation (c_sat_ > 120 μM). Hence, both mutants show much higher c_sat_ compared to WT (c_sat_ ~ 20 μM at 150 mM salt) (Fig. 4B). Notably, 5M→A^IDR^ variant exhibited phase separation at or above 75 mM NaCl, with nearly two-fold higher c_sat_ than WT (Fig. 4A-C). Overall, the observed increase in c_sat_ for 6F→A and 5M→A^IDR^ mutants is consistent with predictions from CG simulations, showing that both of these residues are important for CTD phase separation, despite being outside the CR. However, it is important to note that the effects of methionine residues within the conserved helical domain are substantially greater than those outside the CR, suggesting the dramatic enhancement of phase separation via helix-helix contacts stabilized through methionine residues in the CR.

To test if the importance of methionine residues generalizes to the alphabet of phase separation or only works for five evolutionarily conserved methionine positions within the flanking regions of CTD, 5 alanine residues within CTD flanks are mutated to methionine residues within the 5M→A^CR^ mutant (5A→M^IDR^ + 5M→A^CR^), boosting the total methionine content in the flanking regions from 5 total to 10 total. Interestingly, the resulting mutant has a much lower c_sat_ than 5M→A^CR^ (~70 μM), showing that addition of methionine residues increases phase separation. DIC microscopy images confirm the formation of phase-separated droplets above 70 μM (See Fig. S10). However, in the context of wild-type CR, making the same substitution of alanine to methionine (5A→M^IDR^) results in the formation of non-moving round-shaped clusters at 150 mM NaCl and leads to irregularly shaped amorphous assemblies consistent with aggregation at 300 mM salt (Fig. 4D, S11). Because this variant leads to rapid aggregation, we did not test the phase separation in the saturation concentration assay. The enhanced aggregation hints that the residues in the CTD sequence (especially methionine) are evolutionarily designed for an optimum number of occurrences. These two mutants further highlight the significance of methionine residues in modulating phase separation. A 5M→L^IDR^ mutant showed almost no change compared to WT, whereas 5M→Y^IDR^ lowered the saturation concentration by an order of magnitude (Fig. 3 A-C). These results are consistent with the predictions from the CG simulations, highlighting the importance of aliphatic residues for phase separation along with the aromatic residues.

### Phenylalanine and methionine residues are important for CTD self-assembly

Finally, we evaluated the impact of these mutations inside and outside the CR that affect phase separation on the helical structure of the CR and its helix-helix contacts. NMR spectroscopy is a powerful technique that provides atomic resolution information on biomolecular structure and contacts. The 2D NMR spectrum is the fingerprint of a protein and provides direct information regarding the order/disorder of the protein as well as the formation of stable and transient contacts. Using NMR as a tool, we have previously established that TDP-43 CR alpha-helix self-assembly can be monitored as chemical shift perturbations (CSP) in CR residues upon increasing the protein concentration in conditions where TDP-43 CTD does not readily phase separate (i.e. low salt conditions) (Fig. 5A, WT CTD panel, (Conicella et al., 2016)). To access the impact of mutations that alter TDP-43 CTD phase separation, we compared the 2D NMR spectrum of the variants with that of the WT CTD at 20 μM (a monomeric reference) and 90 μM (Figure 5A). Interestingly, the resonances arising from residue positions in the CR for 5M→A^IDR^ and 6F→A were nearly unperturbed compared to WT CTD at low protein concentration (20 μM), suggesting the alpha-helical structure of the monomeric CR is intact in these constructs. For WT CTD, we observed significant up-field ^15^N CSP at 90 μM for the CR residues corresponding to self-interaction and enhanced helicity (Conicella et al., 2016; Wang and Jardetzky, 2002). However, the CSPs for CR residues in 5M→A^IDR^ were reduced and were minimal in 6F→A compared to WT CTD (Figure 5B), indicating disruption in the helix-helix assembly. Taken together, these data suggest that phenylalanine and methionine outside of the CR form contacts important in helix-mediated CTD self-assembly even in the early stages of assembly preceding phase separation. Importantly, substitution of these residues outside of the CR has no impact CR helicity, consistent with the model that partial CR helical structure is stable and does not require contacts outside the CR.

**Figure 5.**
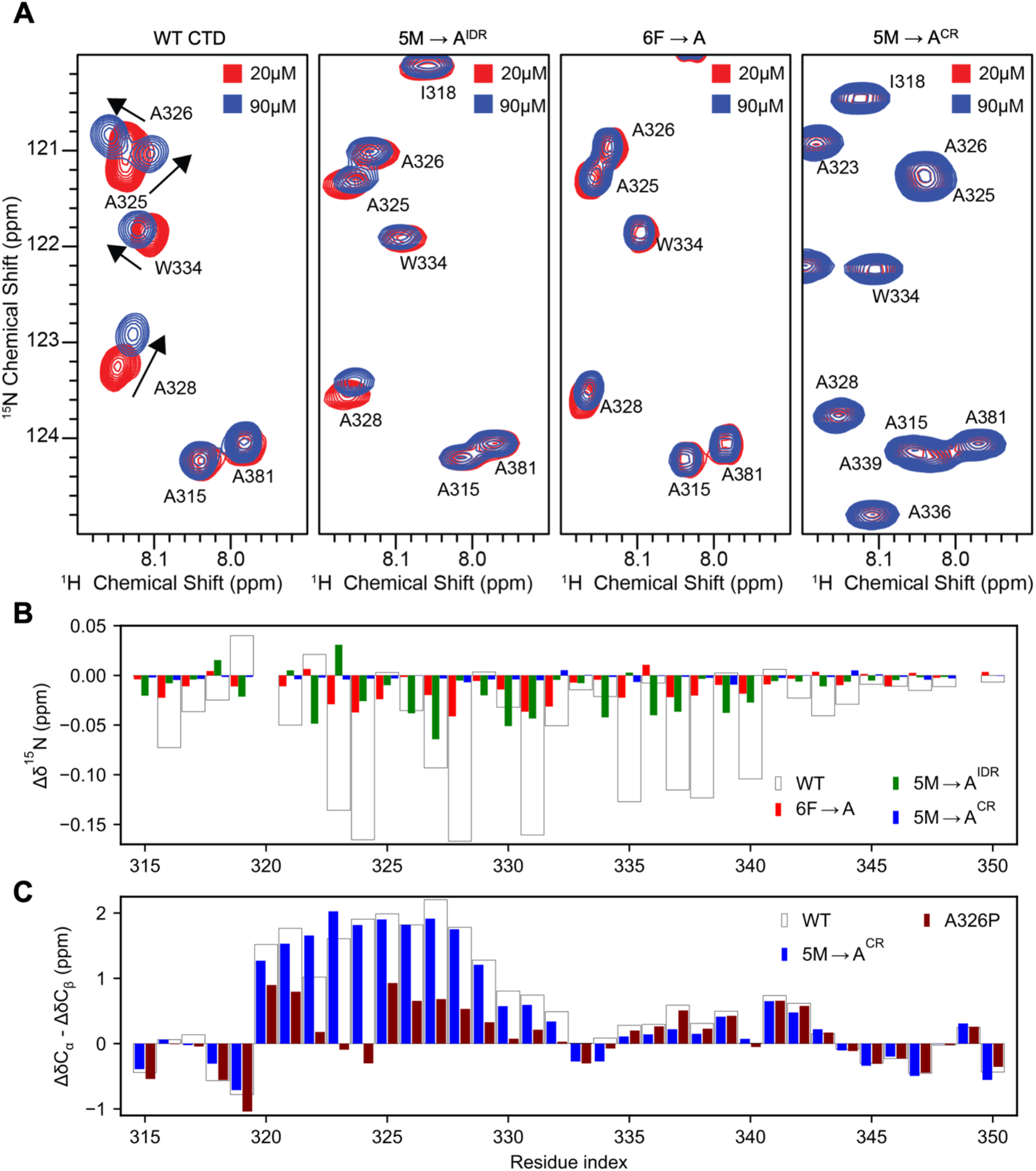
Phenylalanine and methionine residues are crucial for TDP-43 self-assembly. **A**. Overlay of ^1^H-^15^N HSQC spectra of the TDP-43 CTD at two concentrations 20 μM (red) and 90 μM (blue), showing chemical shift perturbations (arrows) associated with intermolecular interactions. The chemical shift deviations are minimal for 6F→A variant and reduced for 5M→A^IDR^ variant, whereas 5M→A^CR^ doesn’t show chemical shift deviations up to 90 μM protein concentrations. **B**. A comparison of ^15^N chemical shift differences (Δδ^15^N) for the CR of WT CTD vs. its mutant variants confirms the disruption of helix-helix interactions by designed variants **C**. Secondary shifts (ΔδCα-ΔδCβ) of 5M→A^CR^ mutant relative to WT CTD and helix-disrupting A326P show that 5M→A^CR^ conserves the helix present in WT CTD

Previously we showed that the helix-disfavoring M337V ALS-associated mutation disrupts helix-helix assembly, as does an engineered proline mutation M337P incompatible with helical structure (Conicella et al., 2016). This observation prompted us to test if the methionine residues play a critical role in helix-helix interactions beyond stabilizing the helix itself. To this end, we tested the 5M→A^CR^ variant (that substitutes all 5 conserved region methionine positions to alanine – M322A, M323A, M336A, M337A, and M339A, see above). As expected, the two-dimensional spectral fingerprint at the CR was significantly perturbed by the substitution of the 5 methionine residues. In order to identify the CR residues, we carried out standard triple resonance NMR assignment experiments (including the HNN experiment, see Methods) and obtained backbone amide resonance assignments. Interestingly, substitution of all 5 conserved region methionine positions to alanine (5M→A^CR^) prevented assembly of the CR region, as little to no CSPs for the CR residues were observed upon increasing concentration (Figure 5A-B). Therefore, these data suggest the methionine residues in the CR are important for mediating helix-helix contacts probably due to loss of methionine side-chain contacts. To test if the mutations disrupt helicity, we calculated the secondary structure content of the CR in 5M→A^CR^. Secondary structure analysis was performed by comparing the experimental C_α_ and C_β_ chemical shifts with their predicted random coil shifts, ΔδC_α_-ΔδC_β_ (Kjaergaard et al., 2011; Kjaergaard and Poulsen, 2011), as we did previously for TDP-43 CTD WT and helix-enhancing mutants (Conicella et al., 2020; Conicella et al., 2016). Surprisingly, our NMR data analysis revealed a significant population of helical structures based on residual ^13^C chemical shifts that are very similar to WT CTD. As a control, these values are also significantly higher than in the helix-disrupting A326P variant (Conicella et al., 2016) (See Fig. 5C). However, it should be pointed out that the secondary structure propensity of alanine residues at 322 and 323 positions are even more positive than those for methionine at these positions in WT CTD, possibly due to the higher helicity for polyalanine sequences (Moreau et al., 2009). Still, the helicity was slightly smaller in other locations compared to WT CR, suggesting that these conserved methionines also form intramolecular contacts to stabilize the helix in TDP-43 CTD. Taken together, these data show that methionines are important for intermolecular contacts that mediate helical assembly and stabilize the helix.

## Discussion

We adopted a hybrid simulation-based strategy coupled with *in vitro* experiments to identify the critical residue types which modulate the phase separation of TDP-43 CTD. Our results indicate that substitution of phenylalanine residues (6F→A) outside CR significantly reduces the phase separation of CTD. Along similar lines, F→S substitution within the CTD was previously shown to disrupt phase separation *in vivo* for a reporter TDP-43 construct and *in vitro* for full-length TDP-43 (Schmidt et al., 2019). In addition, solid-state NMR experiments showed that phenylalanine residues in IDR1/2 were also required for fibril formation from phase-separated droplets, and 6F→A substitution prevented amyloid propagation in cross-seeding experiments (Pantoja-Uceda et al., 2021). Altogether, these observations confirm that phenylalanine residues are critical for phase separation and aggregation of CTD.

TDP-43 CTD contains methionine residues which are equally distributed (five each) in the CR and the flanking regions. We observed that substitution of methionines outside the CR (5M→A^IDR^) significantly weakens phase separation, whereas 5M→A^CR^ mutant disrupts phase separation, highlighting their contribution to phase separation. The “stickiness” of methionine is further confirmed by reversing the reduced phase separation caused by removing the methionines inside the CR (5M→A^CR^) with the addition of methionine residues in the region flanking the CR (5A→M^IDR^ + 5M→A^CR^). These data are consistent with previous efforts showing that MV→A substitutions within the methionine-rich LC domain of Pab-1, which is found to modulate Pab-1 phase-separation, increased the domain’s radius of gyration and demixing temperature (Riback et al., 2017). Additionally, methionines in the FUS RGG3 domain have been shown to make contacts with the FUS SYGQ low complexity domain within a co-condensed phase (Murthy et al., 2021). In sum, methionine residues, surprisingly, substantially contribute to the interactions that drive the phase separation of CTD.

It is important to consider the relative contribution to phase separation of methionine compared to other aromatic and aliphatic residues. CG simulations and experimental data show that it has lower stabilizing contributions to phase separation than aromatic residue – tyrosine (5M→Y^IDR^). Substitutions of methionine with another aliphatic residue, leucine (5M→L^IDR^), yielded minor changes in phase separation compared to WT by experiment and simulation. Likewise, 5M→S mutation in IDR1/2 is shown to increase TDP-43 reporter assay droplet dynamics without affecting its c_sat_, while 5M→V behaved similarly to WT under *in vivo* conditions (Schmidt et al., 2019). Our observations here provided new direct evidence for the contribution of methionine to the thermodynamics of phase separation and not just the dynamics of the condensed phase. Furthermore, our results suggest that other aliphatic residues except alanine may have similar contributions, expanding the molecular grammar of low complexity domain phase separation.

Our sequence alignment analysis over 93 homologs of TDP-43 CTD (Schmidt et al., 2019) shows that the number of aliphatic residues (L=2, I=1, and M=5) outside CR is mostly conserved and switched to another aliphatic residue (VLIM) in some homologs (e.g., L270, I383) (See Fig. S12). On the other hand, alanine, which is also aliphatic, is less conserved and tends to switch to or appear as a replacement for polar residues. Correspondingly, our *in vitro* experiments support the idea that alanine residues outside CR contribute less to LLPS than larger aliphatic residues.

Although aliphatic residues may contribute to phase separation similarly based their similar hydrophobicity and size, it is important to note that methionine has unique physicochemical properties that could broaden its role in phase separation (Aledo, 2019). Foremost, it has highly flexible, unbranched side chains that could provide additional conformational dynamics compared to other hydrophobic residues and enable important contacts required for intermolecular self-assembly (Gellman, 1991). Recently, Neuweiler and coworkers demonstrated that methionine side chains in the spidroin NTD core promote dimerization, which was dramatically impaired by the simultaneous mutations of all methionine to leucine, despite yielding higher stability and fully preserving its structure (Heiby et al., 2019). Our results suggests that the structural plasticity of methionine sidechains could contribute to the TDP-43 CTD helix-helix contacts and higher-order assembly in a similar manner, which explains the abolishment of phase separation via 5M→A^CR^ mutant even though it preserves the helicity.

Another important characteristic of methionine is the reversible oxidation (Achilli et al., 2015; Moskovitz et al., 1997) of solvent-exposed methionine sidechains to methionine sulfoxide (MetO) which allows the regulation of the protein phase separation. This was probed in detail by the recent studies from the Benjamin Tu (Yang et al., 2019) and Steven McKnight (Kato et al., 2019) laboratories wherein the methionine-rich LC domain self-association of yeast ataxin-2 was demonstrated to be controlled by reversible methionine oxidation. Similarly, phase separation of TDP-43 CTD was shown to be disrupted *in vitro* due to the non-specific oxidation of methionines in the presence of H_2_O_2_, an oxidizing agent, and restored via enzymatic reduction of methionine oxidation (Lin et al., 2020). However, those previous results do not provide direct evidence of why methionine oxidation disrupts phase separation. Our findings comparing methionine to alanine by explicit evaluation of the saturation concentration directly attribute methionine’s role in phase separation to favorable methionine residue contacts.

To conclude, our study provides new evidence that a delicate balance of many residue types tunes the phase behavior of TDP-43 CTD. In particular, phenylalanine and methionine residues across both the disordered and helical regions significantly influence the CTD phase separation. Our NMR spectroscopy analysis suggests that phenylalanine and methionine residues are significant for TDP-43 intermolecular contacts and that substitution of these positions decrease TDP-43 assembly. Interestingly, the cryo-EM structure (Arseni et al., 2022) of aggregated TDP-43 from the human brain of an individual who had ALS with FTLD has a cluster of methionine and phenylalanine residues in the core (M307, M311, F313, M322, M323) and numerous methionine-aromatic contacts can be seen at different locations. This implies the possible presence of energetically stabilizing aliphatic-aromatic contacts that facilitates the TDP-43 self-assembly and may promote fibrillization as well.

The results reported herein show the importance of methionine residues in TDP-43 CTD phase separation and could be general for other PrLDs. Upon methionine composition analysis among 29 human RBP prion candidates (King et al., 2012), including ataxin-2, we found that one-third of those PrLDs contain at least 4% methionine (Table S2, Fig. S13). The majority of those RBPs are shown to phase-separate either by increasing protein concentration or using a molecular crowding agent (Wang et al., 2018b). Based on these data, we suspect that methionine residues may also have contributed to evolution by promoting the assembly of biomolecular condensates. Finally, our findings could help us to paint a more accurate picture of the phase separation of RBPs.

## Material and Methods

### All-atom PT-WTE simulation protocol

PT-WTE simulations of TDP-43 CTD (aa:267-414) were performed using GROMACS 2018.8 (Abraham et al., 2015) and PLUMED 2.5.4 (Tribello et al., 2014). PT-WTE requires significantly fewer temperature replicas compared to conventional parallel tempering due to the amplification of potential energy fluctuations through periodic deposition of gaussian potentials (Deighan et al., 2012). A total of 20 temperature replicas were generated based on a geometric progression that spanned a range from 300 to 475 K. The system topology was modeled using the AMBER99SBws-STQ force field (Best and Mittal, 2010; Best et al., 2014; Tang et al., 2020). Each replica was simulated for ~585 ns, resulting in an aggregate simulation time of ~12 μs.

The initial structure for PT-WTE simulation was obtained from a CG simulation (1 μs) of CTD (aa:267-414) using the HPS-Urry CG model (Dignon et al., 2018b; Regy et al., 2021) with helical conformations in aa: 320-341 imposed using pseudo-dihedral potentials. The CG model was then converted into an atomistic model using MODELLER (Sali and Blundell, 1993) and solvated using TIP4P2005 (Abascal and Vega, 2005) water molecules in an octahedral box (l=12.5 nm). 100 mM NaCl was added to mimic physiological salt concentration, and three additional Cl^−^ ions were added to achieve electroneutrality. Following energy minimization, temperature, and pressure equilibration of the system at 300 K and 1 bar, temperature replicas were prepared and equilibrated without exchange for 10 ns at higher temperatures using the Langevin integrator (τ_t_= 1 ps). Following temperature equilibration of the replicas, production runs (timestep = 2 fs) were performed with bias potentials deposited every 4 ps and exchanges attempted between adjacent replicas every 1 ps. The initial height and width of the gaussian potential was set to 1.5 and 450.0 kJ mol^−1^ respectively. Exchange probabilities between adjacent replicas varied from 20-40%.

The first 85 ns from the production replicas were discarded as equilibration time, during which the initial height of the gaussian potential (deposited every 4 ps) dropped below 0.25 kJ mol^−1^ (Fig. S2A), and sufficient overlap in the potential energy distributions (Fig. S2B) between adjacent temperature replicas was observed. Both 300 and 346.5 K replicas underwent several round trips (Fig. S2C) through the entire temperature space on the ~100 ns timescale, which indicates efficient conformational exchange between replicas. The convergence of global properties was determined based on the radius of gyration (R_g_) and end-to-end distance computed for the 300 K ensemble. The low errors associated with their probability distributions averaged over 100 ns intervals indicate that the ensemble is well-converged (Fig. S2D). Per-residue interaction energies were computed using gRINN (Sercinoglu and Ozbek, 2018).

### Coarse-grained simulation protocol and analysis

Coexistence phase simulations of TDP-43 CTD wild type and mutants were conducted using the HOOMD-Blue 2.9.3 software package (Anderson et al., 2020), according to the protocol described in our previous studies (Mammen Regy et al., 2021). Residue beads from the conserved helix (aa:320-341) domain were kept rigid during the CG simulations to mimic the α-helical structure present in CR. The initial slab configuration (15 nm × 15nm × 1050 nm) was prepared from 100 chains of the TDP-43 CTD sequence based on the HPS-Urry model (Regy et al., 2021) and simulated for 5 μs in the NVT ensemble at 300 K using a time step = 10 fs. The Langevin thermostat was used for temperature control, with the damping factor (τ) set to 1000 ps. Under the same simulation protocol, CG single chain simulations of wild type TDP-43 CTD were conducted at 300 K using the LAMMPS software package (Thompson et al., 2022). During all analysis, the first 1 μs was treated as an equilibration period and omitted for subsequent calculations. Radius of gyration, coexistence densities and pairwise contact probabilities were calculated using the protocol mentioned in our previous work (Dignon et al., 2018b; Mammen Regy et al., 2021).

### Expression and purification of recombinant proteins

Wild-type human TDP-43 CTD and mutant variants of TDP-43 CTD were expressed from a codon-optimized sequences in the pJ411 bacterial expression vector. These proteins were expressed in BL21 Star (DE3) Escherichia coli cells (Life Technologies). The expression and purifications were either in LB or M9 minimal media supplemented with ^15^NH_4_Cl with slight modifications as previously described (Conicella et al., 2020; Conicella et al., 2016). Briefly, the bacterial cultures were induced at OD 0.8 with 1 mM IPTG and grown for 4h at 37°C and 200 rpm. The cells were harvested by centrifugation (6000 rpm, 15 minutes, 4°C). Cell pellets thus obtained were resuspended in 20 mL buffer (20 mM Tris, 500 mM NaCl, 10 mM imidazole, pH 8.0), lysed with an ultrasonic cell disrupter, and centrifuged (20,000 g, 1 hr, 4°C). The insoluble material with the inclusion bodies was resuspended in 40 ml solubilizing buffer (8M urea, 20 mM Tris, 500 mM NaCl, 10 mM imidazole, pH 8.0), and centrifuged (25,000 rpm, 1hr, 19°C). The supernatant was filtered by 0.22 μm syringe filter, and the protein was purified using a 5 mL Histrap HP column with a gradient of 10 to 500 mM imidazole added to the insoluble buffer. The pure protein fractions were desalted using a HiPrep 26/60 Desalting Column into TEV cleavage buffer (20 mM Tris, 500 mM GdnHCl, pH 8.0) for TEV reaction overnight at room temperature. After cleavage, the solid urea was added to the solution to approximately 8 M urea and the resolubilized protein was applied to the Histrap HP column to remove histidine-tag and histidine-tagged TEV protease. Cleaved protein fractions were concentrated and buffer-exchanged into storage buffer (20 mM MES, 8 M urea, pH 6.1), flash-frozen, and stored at −80°C.

### Microscopy

According to the manufacturer’s instructions, using 0.5 mL Zeba spin desalting columns (Thermo Scientific), the protein samples were desalted to MES buffer (20 mM MES, pH 6.1), and diluted to 40 μM, supplemented with varying concentrations of 0, 150, and 300 mM NaCl. Samples were gently mixed and monitored for phase separation in DIC micrographs with a Zeiss Axiovert 200M microscope with a 40x objective after spotting 10 μL of sample onto the coverslip. The images were processed with ImageJ.

### Determination of saturation concentration for quantification of *in vitro* phase separation

For a quantitative analysis of CTD mutant phase separation, we conducted assays to determine the saturation concentration wherein the protein concentration in the supernatant was measured after centrifugation of samples with increasing salt concentration to induce phase separation. During centrifugation, the micrometer-sized droplets sediment account for protein in the phase-separated state, while the sample remaining in the supernatant accounts for the protein content in the dispersed phase. The amount of protein remaining in the supernatant represents the saturation concentration (the concentration above which the protein phase separates) and gradually reduces with increasing salt concentrations and is correlated with phase separation propensity. The desalted protein samples were diluted to 80 μM or 120 μM using the MES buffer. Phase separation was initiated by adding an equal volume of NaCl stock solution prepared in MES buffer to obtain final salt concentrations of 0, 75, 150, 300, and 500 mM NaCl. Then, the samples were gently mixed and centrifuged for 10 minutes at 12,000 rpm. Later, the protein concentration in the supernatant was measured using the (A280 E280 = 18,350 M^−1^ cm^−1^) with a Nanodrop 2000c. All data points were measured in triplicate.

### NMR spectroscopy

All NMR experiments were performed Bruker Avance 850 MHz or 600 MHz ^1^H Larmor frequency spectrometers with HCN TCI z-gradient cryoprobes at 298K. The NMR samples were prepared in 20 mM MES Buffer, pH 6.1, supplemented with 10% D_2_O. Backbone assignments for the CTD variant were obtained by standard triple resonance experiments (HNCACB, HNCA, CBCACONH, and HNN (HNCANNH)). The NMR data were processed with Bruker Topspin and analyzed using CCPNMR (Skinner et al.). Experimental NMR parameters summary can be found from our previous works (Conicella et al., 2020; Conicella et al., 2016).

## Data and resource availability

NMR chemical shift assignments in this paper for TDP-43 CTD WT (26823), A326P (26828), 6F→A, 5M→A^IDR^, 5M→A^CR^ (deposition in process for new variants) can be obtained online from the Biological Magnetic Resonance Database (BMRB, http://www.bmrb.wisc.edu/). Plasmids generated for this project can be found at Addgene.org (deposition in process). Associated protocols and simulation code can be obtained by contacting the corresponding authors.

## Acknowledgments

P.M. thanks Nina Jovic for helping with setting up PT-WTE simulations. This work was supported by NINDS and NIA R01NS116176. Atomistic and coarse-grained simulations were conducted with the advanced computing resources provided by Texas A&M High Performance Research Computing.

## Competing interests

The authors declare that they have no competing interests.

## Notes

### Competing Interest Statement

The authors have declared no competing interest.

