## Supplementary Information for "Aliphatic residues contribute significantly to the phase separation of TDP-43 C-terminal domain"

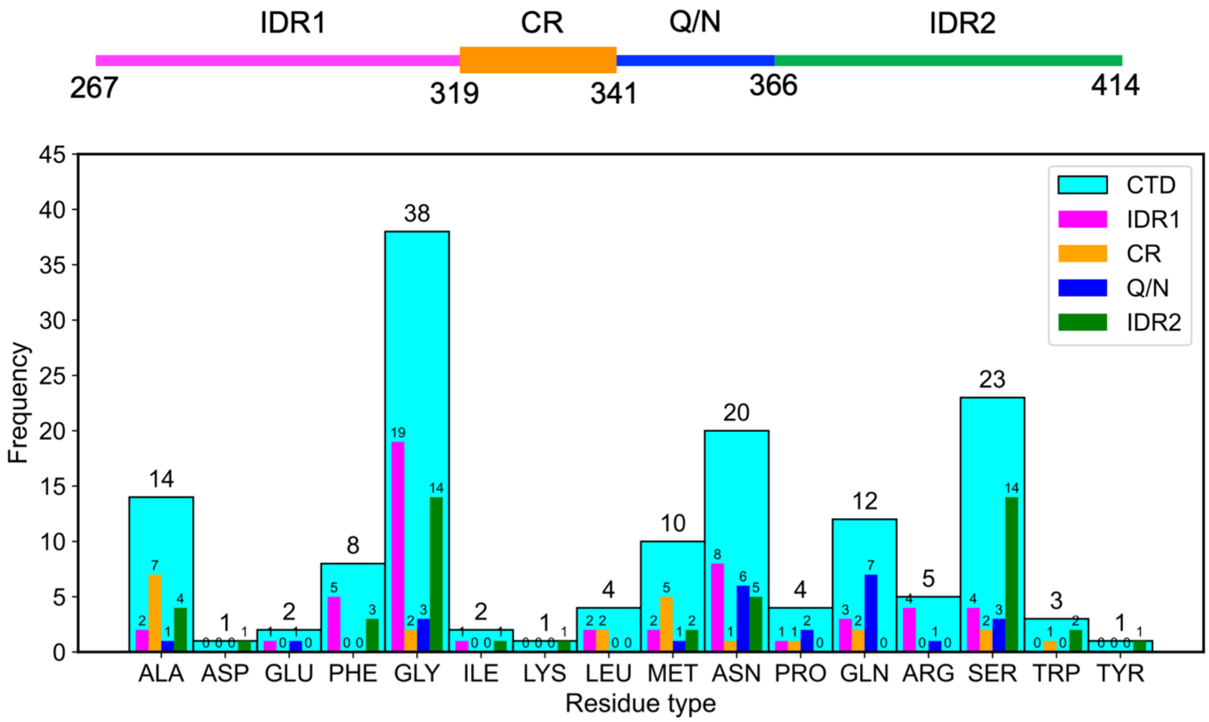


**Figure S1.** Sequence Composition of TDP-43 C-terminal domain (CTD) and its subdomains

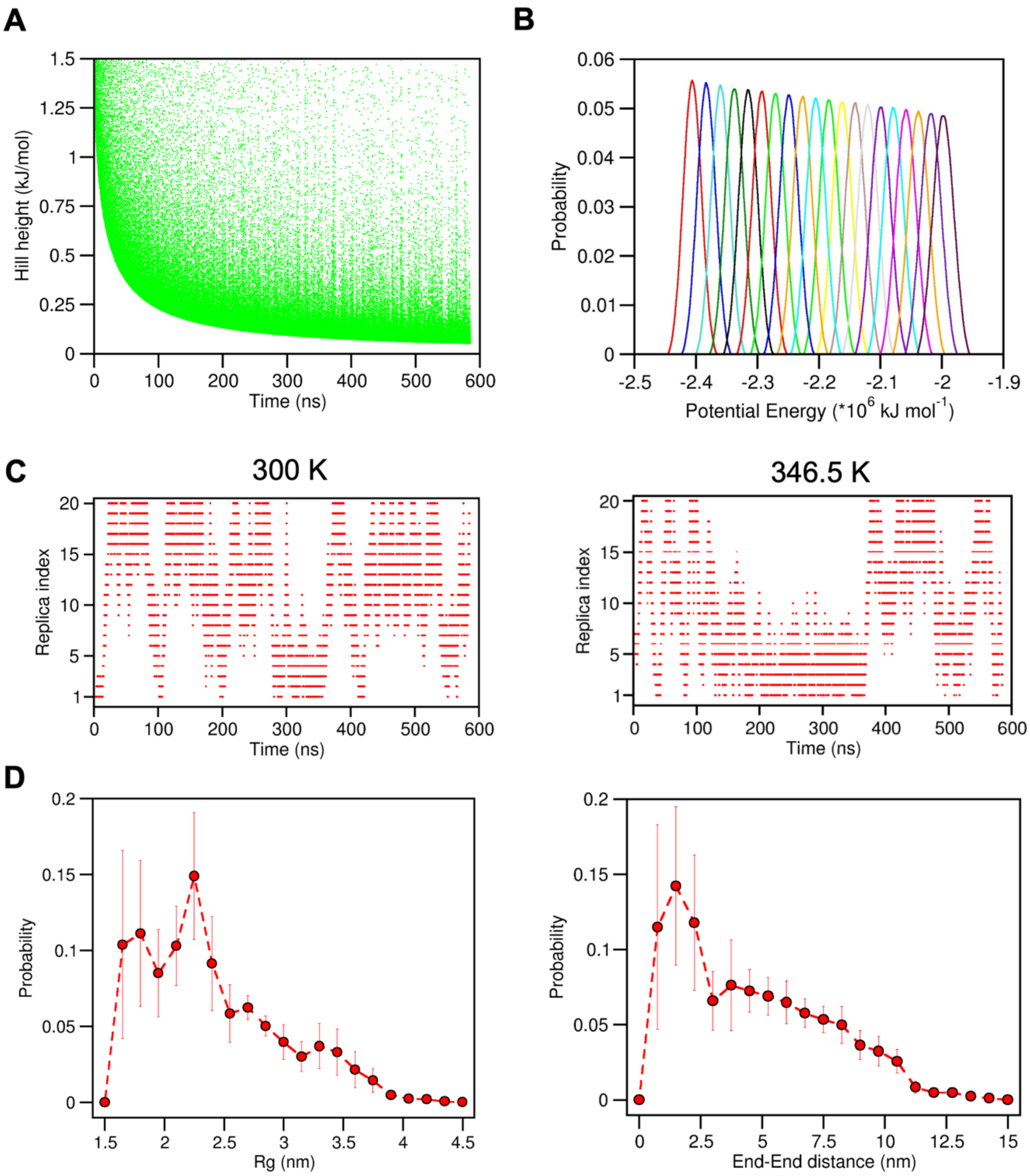


**Figure S2.** Assessing convergence of the PT-WTE ensemble. **A.** Variation of gaussian hill height as a function of time for the 300 K replica. **B.** Potential energy distributions for all 20 replicas. Distributions were calculated over ~585 ns per replica. **C.** Movement of 300 K and 346.5 K replicas through temperature space. **D.** Radius of gyration and end-end distance probability distributions of the 300 K replica calculated over 500 ns. The first ~85 ns was excluded as equilibration time.

**A**

| Chemical shift | RMSD (ppm) |
|---|---|
| C’ | 0.42 |
| $C_\alpha$ | 0.39 |
| $C_\beta$ | 0.38 |

**B**

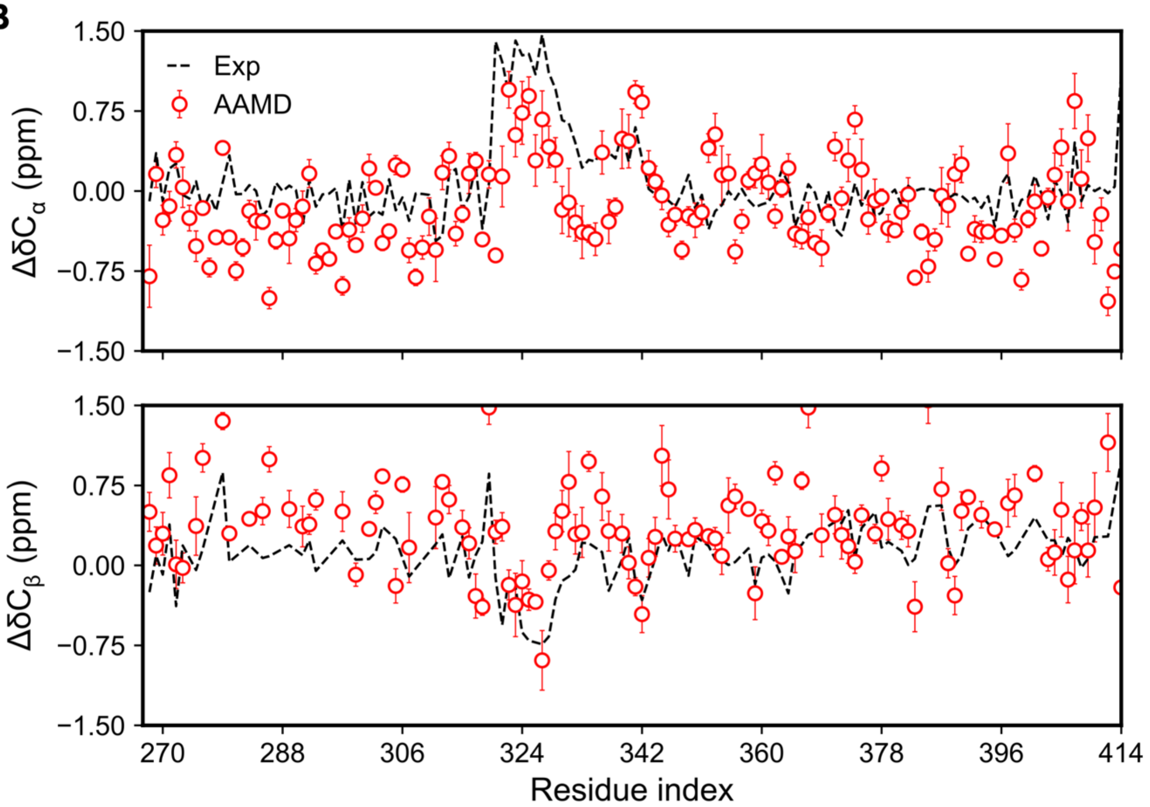


**Figure S3.** Validation of CTD ensemble at 300 K based on NMR observables. **A.** Chemical shift RMSDs between the 300K replica and experiment. **B.** Comparison of per-residue secondary chemical shifts (Cα, Cβ) between 300 K simulation replica and experiment

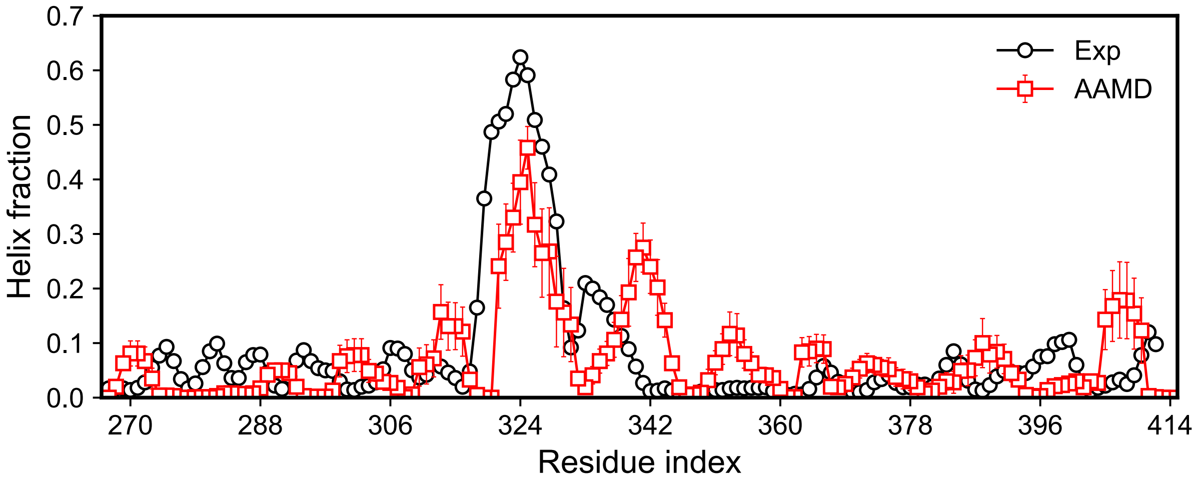


**Figure S4.** Comparison of per-residue helical fractions from simulation with NMR-based fractions calculated using δ2D (Camilloni et al., 2012)

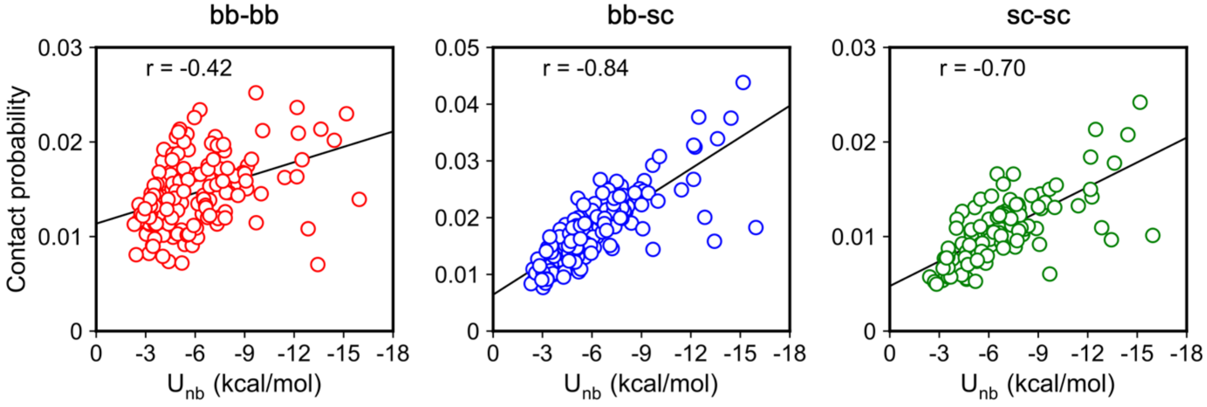


**Figure S5.** Correlation between per-residue intramolecular contacts and interaction energies (coulombic + van der Waals) for neutral residues calculated from single-chain atomistic simulations.

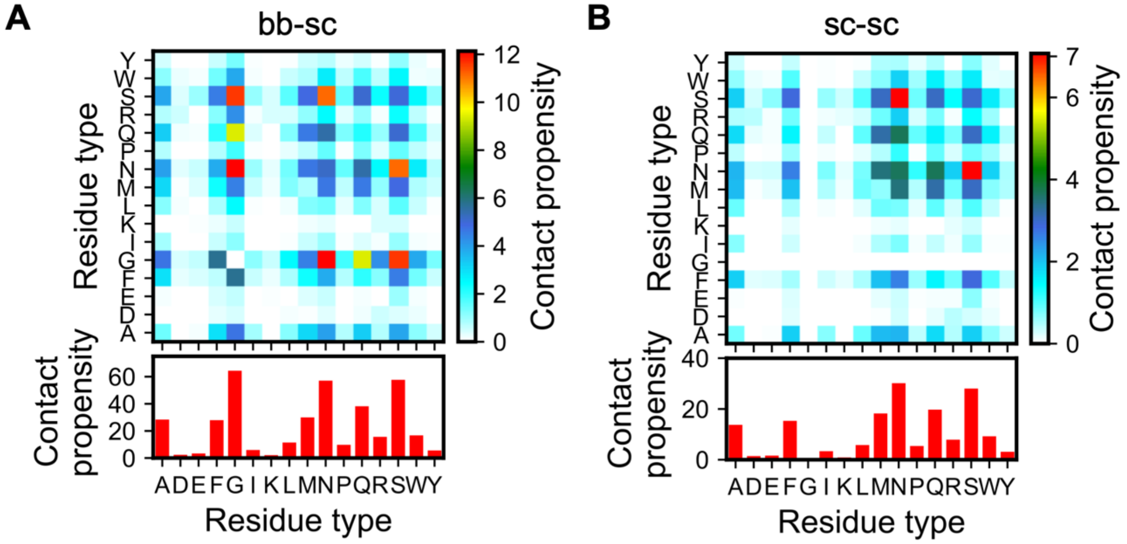


**Figure S6.** Pairwise interactions between all residue types occurring in CTD decomposed into bb-sc and sc-sc interaction modes are shown in A and B, respectively. The bottom panels show the one dimensional summation.

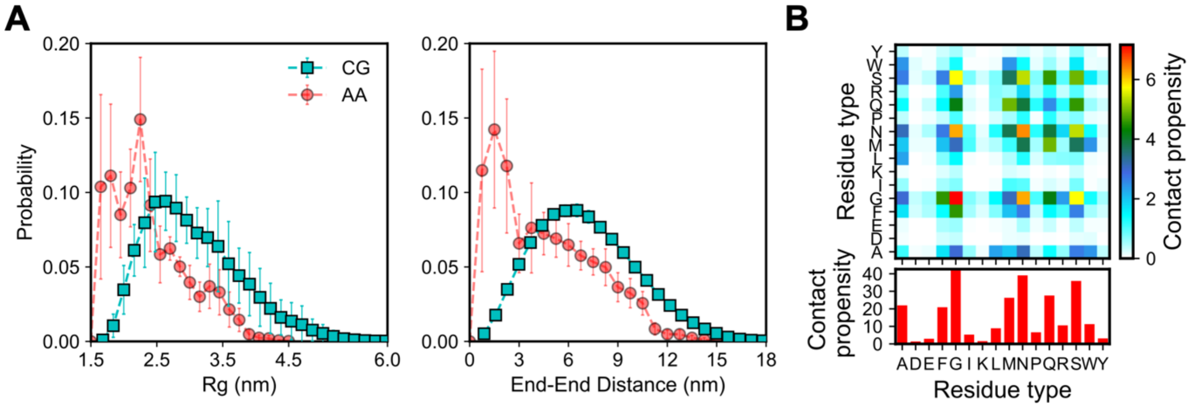


**Figure S7. A.** Radius of gyration and end-end distance probability distributions of the single chain TDP-43 CTD from CG simulations calculated over 5 μs. The first 1 μs was excluded as equilibration time. Errors are estimated using block averages with four blocks. **B.** Intramolecular pairwise interactions between all residue types occurring in single-chain CTD from CG single-chain simulations are in good agreement with those for atomistic simulations.

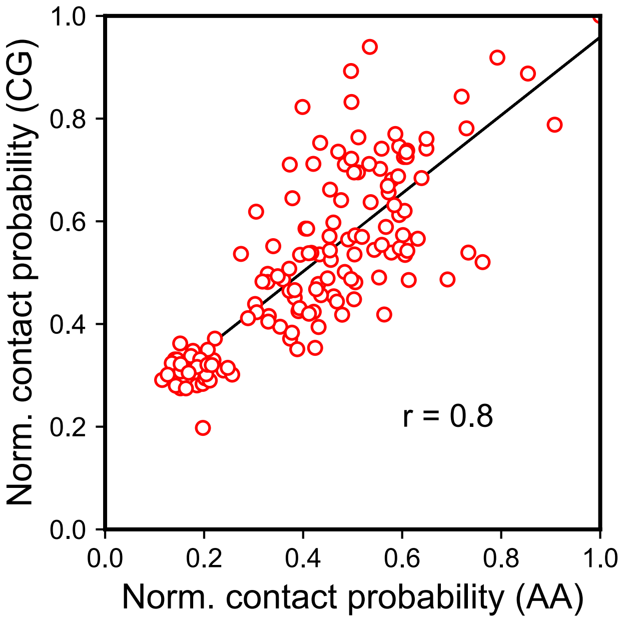


**Figure S8.** The correlation between per-residue intramolecular and intermolecular contact probabilities is calculated from all-atom single-chain simulations (AA), and coarse-grained (CG) condensed phase simulations, respectively. The probabilities were normalized by the maximum value observed for a given residue in each case.

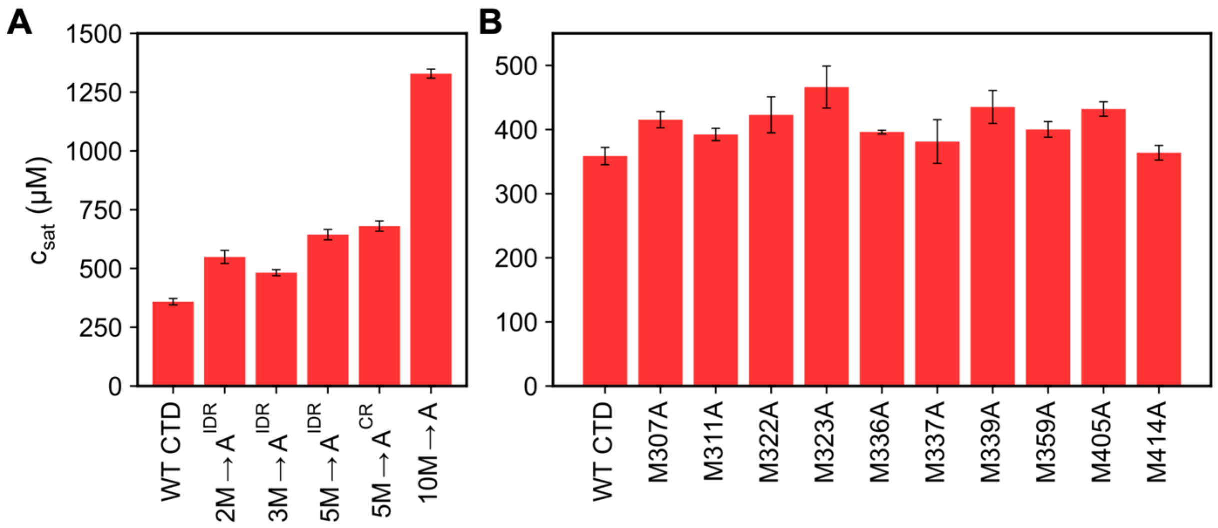


**Figure S9.** Saturation concentration ($c_{sat}$) of bulk and positional methionine mutations for TDP-43 CTD computed from CG phase coexistence simulations.

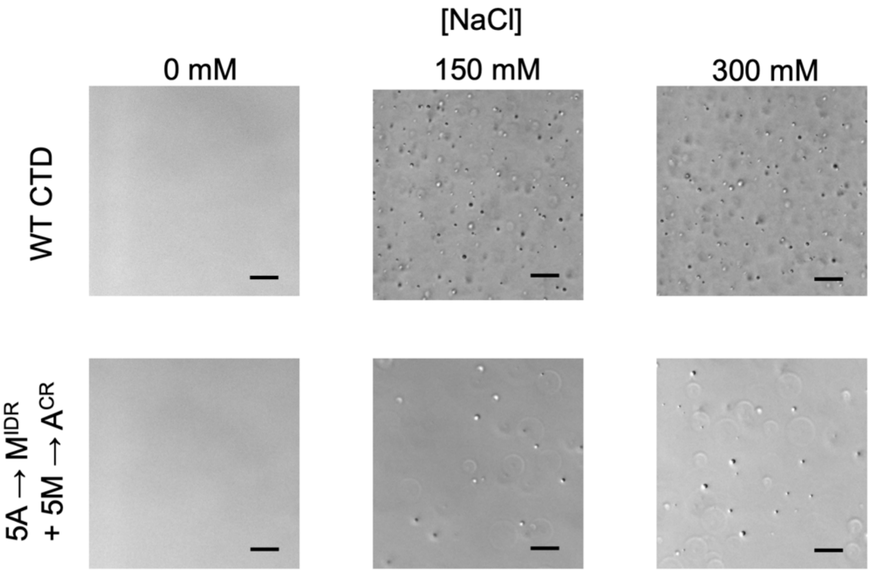


**Figure S10.** DIC micrographs for WT TDP-43 CTD and A→M$^{IDR}$ + 5M→A$^{CR}$ variant at different salt concentrations and 80 μM protein concentration. Scale bar, 20 μm.

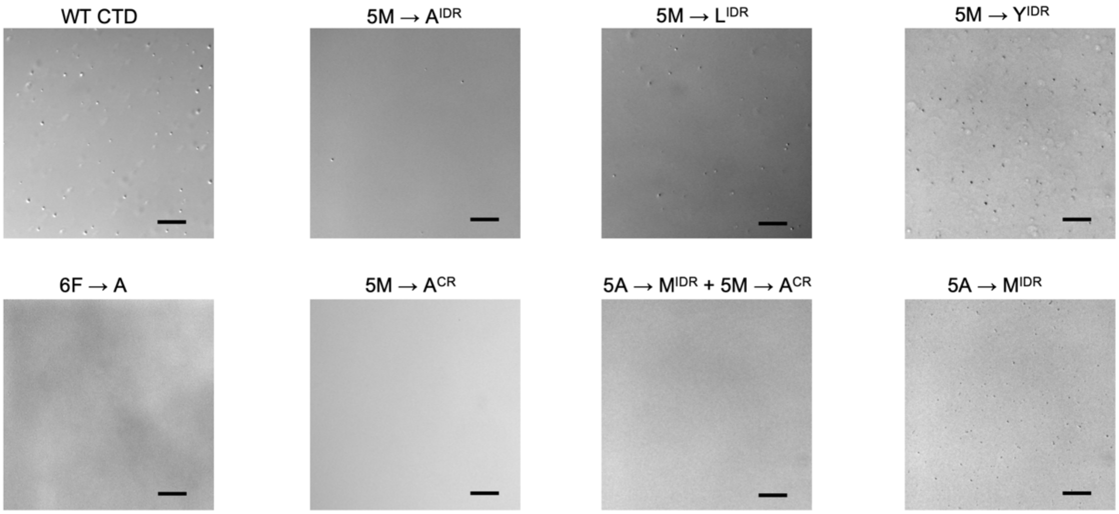


**Figure S11.** DIC micrographs for WT TDP-43 CTD and its designed variants at 150 mM salt and 20 μM protein concentration. Scale bar, 20 μm.

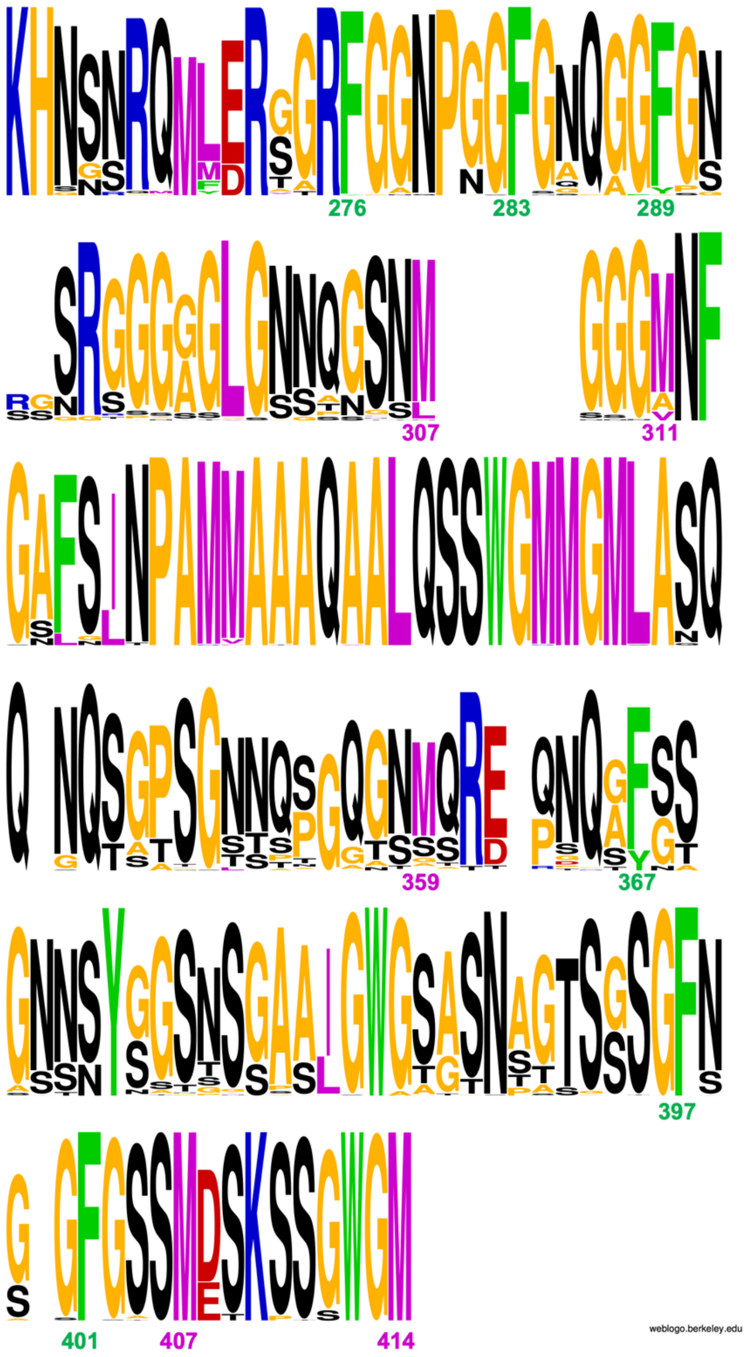


**Figure S12.** The sequence logo of TDP-43 CTD is based on the sequence alignment of 93 vertebrate homologs (Schmidt et al., 2019). Sequence alignment was carried out using MEGA11 based on the MUSCLE option (Tamura et al., 2021). Sequence logos were produced by using WebLogo 3 (Crooks et al., 2004). The gap between M307 and G308 comes from Xenopus laevis CTD homolog that has additional residues (GSGGGG) between M307 and G308 present in homo sapiens CTD.

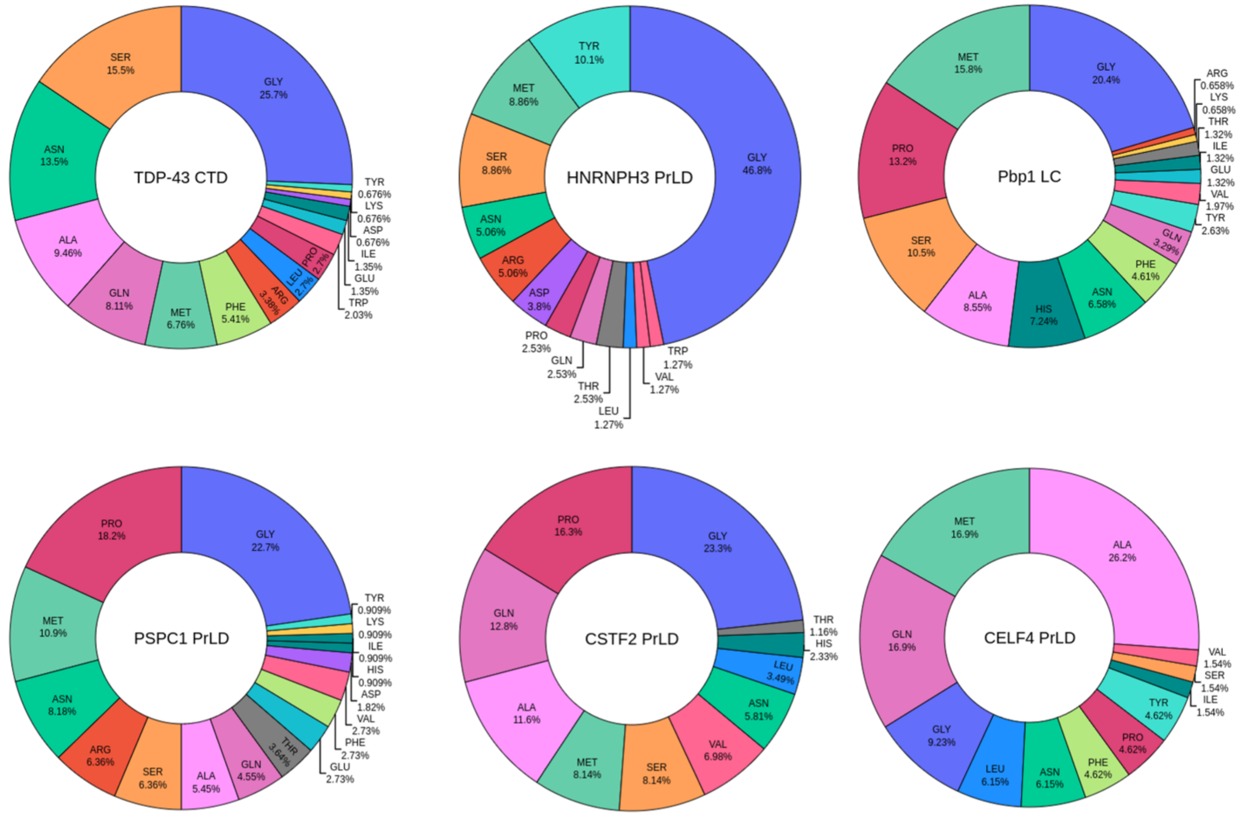


**Figure S13.** Residue composition of human RNA-binding proteins with prion-like domains (PrLDs) that have significant amount of methionine composition.

**Table S1.** TDP-43 CTD mutants were tested in CG condensed phase simulations and *in vitro* assays.

| Mutant | Residues |
|---|---|
| 6F→A | F276A, F283A, F289A, F367A, F397A, F401A |
| 5M→A$^{CR}$ | M322A, M323A, M336A, M337A, M339A |
| 5M→X$^{IDR}$ | M307X, M311X, M359X, M405X, M414X, where X = A, L, Y |
| 5A→M$^{IDR}$ + 5M→A$^{CR}$ | A297M, A367M, A381M, A388M, A391M<br>M322A, M323A, M336A, M337A, M339A |
| 5A→M$^{IDR}$ | A297M, A367M, A381M, A388M, A391M |

**Table S2:** Methionine composition analysis of the 29 human RNA-binding proteins with prion-like domains (PrLDs) (King et al., 2012), including ataxin-2, show that one-third of PrLDs is comprised of more than 4% MET.

| PrLD name | **% MET** | **# MET** |
|---|---|---|
| Pbp1 LC (ataxin-2) | 15.79 | 24 |
| CELF4 PrLD | 16.92 | 11 |
| PSPC1 PrLD | 10.91 | 12 |
| HNRNPH3 PrLD | 8.86 | 7 |
| CSTF2 PrLD | 8.14 | 7 |
| **TDP-43 CTD** | **6.76** | **10** |
| HNRNPH1 PrLD | 5.88 | 4 |
| HNRNPH2 PrLD | 5.88 | 4 |
| RBM33 PrLD | 4.27 | 5 |
| TIA1 PrLD | 4.21 | 4 |
